# Antimicrobial and antibiofilm activity of human recombinant H1 histones against bacterial infections

**DOI:** 10.1101/2024.04.03.587932

**Authors:** Betsy Veronica Arévalo-Jaimes, Mónica Salinas-Pena, Inmaculada Ponte, Albert Jordan, Alicia Roque, Eduard Torrents

## Abstract

Histones possess significant antimicrobial potential, yet their activity against biofilms remains underexplored. Moreover, concerns regarding adverse effects limit their clinical implementation. We investigated the antibacterial efficacy of human recombinant histone H1 subtypes against *Pseudomonas aeruginosa* PAO1, both planktonic and in biofilms. After the *in vitro* tests, toxicity and efficacy were assessed in a *P. aeruginosa* PAO1 infection model using *Galleria mellonella* larvae. Histones were also evaluated in combination with ciprofloxacin and gentamicin. Our results demonstrate antimicrobial activity against of all three histones against *P. aeruginosa* PAO1, with H1.0 and H1.4 showing efficacy at lower concentrations. The bactericidal effect was associated with a mechanism of membrane disruption. *In vitro* studies using static and dynamic models showed that H1.4 had antibiofilm potential by reducing cell biomass. Neither H1.0 nor H1.4 showed toxicity in *G. mellonella* larvae, and both increased larvae survival when infected with *P. aeruginosa* PAO1. Although *in vitro* synergism was observed between ciprofloxacin and H1.0, no improvement over the antibiotic alone was noted *in vivo*. Differences in antibacterial and antibiofilm activity were attributed to sequence and structural variations among histone subtypes. Moreover, the efficacy of H1.0 and H1.4 was influenced by the presence and strength of the extracellular matrix. These findings suggest histones hold promise for combating acute and chronic infections caused by pathogens such as *P. aeruginosa*.

**Importance:** The constant increase of multidrug-resistant bacteria is a critical global concern. The inefficacy of current therapies to treat bacterial infections is attributed to multiple mechanisms of resistance, including the capacity to form biofilms. Therefore, the identification of novel and safe therapeutic strategies is imperative. This study confirms the antimicrobial potential of three histone H1 subtypes against both Gram-negative and Gram-positive bacteria. Furthermore, histones H1.0 and H1.4 demonstrated *in vivo* efficacy without associated toxicity in an acute infection model of *Pseudomonas aeruginosa* PAO1 in *Galleria mellonella* larvae. The bactericidal effect of these proteins also resulted in reduction in biomass of *P. aeruginosa* PAO1 biofilms. Given the clinical significance of this opportunistic pathogen, our research provides a comprehensive initial evaluation of the efficacy, toxicity, and mechanism of action of a potential new therapeutic approach against acute and chronic bacterial infections.

## Introduction

Bacterial infections caused by multidrug-resistant bacteria are increasingly burdening the global healthcare system (1, 2). Bacteria have evolved various mechanisms to counteract the harmful effects of antibiotic molecules, including restriction drug uptake, upregulating efflux pumps, altering drug targets, and inactivating drugs (2). Additionally, bacteria have developed defense strategies at the multicellular level. Bacterial biofilms are communities of bacteria growing together and working collectively to withstand threats that would kill planktonic cells (3). Cells in biofilms produce an extracellular matrix (ECM) that surrounds and protects bacteria against antimicrobials, the immune system, phages, and other attacks (3). The biofilm structure and ECM create diffusion gradients of nutrients and antimicrobial molecules, promoting metabolic adaptations that result in population heterogeneity, adding more complexity to biofilm eradication (3, 4). Consequently, biofilm-associated infections are clinically challenging and often become persistent and difficult to treat (5).

*Pseudomonas aeruginosa* is one of the six highly drug-resistant bacteria worldwide (2). Its metabolic adaptability, coupled with an arsenal of virulence factors, explains its prevalence in life-threatening acute and chronic infections, particularly among immunocompromised patients (6). Its ability to form biofilms with increased tolerance to antibiotics leads to the development of chronic wounds and pulmonary failure in patients with cystic fibrosis and other respiratory diseases (6, 7).

The urgent need to develop new antibiofilm strategies that can either replace or complement current alternatives has led to the identification of effective natural compounds, including phytochemicals, biosurfactants, and antimicrobial peptides (AMPs) (5). AMPs are an important part of the innate immunity of multiple species, and acquiring resistance to them is more complicated than to antibiotics (4, 8).

Histones are crucial proteins in eukaryotic DNA organization and regulation (9). However, they can be secreted passively (necrosis) or actively (apoptosis and neutrophil NETosis) into the extracellular fluid (9, 10). Their cationic characteristics lead to cell membrane damage upon interacting with phospholipids (9). Like AMPs, the antimicrobial potential of histones has been demonstrated both *in vitro* and *in vivo* by effectively killing bacteria, fungi, parasites, and viruses (10, 11).

Histone H1 is a basic protein that regulates the higher-order structure of chromatin. It has three structural domains: the N-terminal domain (NTD), the globular domain (GD), and the C-terminal domain (CTD). The CTD represents approximately half of the protein and has the highest content of basic residues (12). Human somatic cells can express up to seven H1 subtypes: H1.0-H1.5 and H1X (13, 14). Histone H1 was proposed as a component of the antimicrobial defense in the human gastrointestinal tract in the 90s (15, 16). Since then, its antimicrobial activity has been reported against various bacteria (10, 11, 17), however, its potential as antibiofilm therapy and *in vivo* efficacy remains largely unexplored.

This study aims to evaluate the activity of three human recombinant histone H1 subtypes against *P. aeruginosa* PAO1 planktonic and biofilms growing cells. We selected two of the more abundant subtypes in mammalian cells, H1.2 and H1.4, and the subtype that increases in differentiated cells, H1.0 (Full length and CTD). Furthermore, we evaluated their potential as combined treatment with antibiotics (Ciprofloxacin and Gentamicin) and assessed their toxicity and efficacy *in vivo* using the animal model *Galleria mellonella*.

## Results

### Antimicrobial efficacy and synergy assessment

The antimicrobial efficacy of histones H1.0, H1-0 C-terminal domain (CTD), H1.2 and H1.4 was determined by measuring their minimal inhibitory concentration 50% (MIC_50_) (18) in the bacterial growth of three different important pathogens (Table 1). Overall, higher antimicrobial activity was observed against both Gram-negative bacteria, *P. aeruginosa* PAO1 and *E. coli* CFT073, with the first being more susceptible. Conversely, a minor effect was observed against the Gram-positive bacteria *S. aureus*.

**TABLE 1.** Antibacterial and cytotoxic activity of the different histones.

|  | MIC<sub>50</sub><sup>a</sup> |  |  | *In vivo*<br>LD<sub>50</sub><sup>b</sup> |
| --- | --- | --- | --- | --- |
|  | *P. aeruginosa*<br>PAO1 | *E. coli* CFT073 | *S. aureus* |  |
| **H1.0** | 55 µg/mL<br>(2.5 µM) | 154 µg/mL<br>(7 µM) | >220 µg/mL<sup>c</sup><br>(>10 µM) | >20 mg/Kg<sup>d</sup><br>(>400 µg/mL) |
| **H1.0 C-ter** | 58 µg/mL<br>(5 µM) | >116 µg/mL<sup>c</sup><br>(>10 µM) | >116 µg/mL<sup>c</sup><br>(>10 µM) | >20 mg/Kg<sup>d</sup><br>(>400 µg/mL) |
| **H1.2** | 111 µg/mL<br>(5 µM) | 222 µg/mL<br>(10 µM) | >222 µg/mL<sup>c</sup><br>(>10 µM) | >20 mg/Kg<sup>d</sup><br>(>400 µg/mL) |
| **H1.4** | 46 µg/mL<br>(2 µM) | 98 µg/mL<br>(4 µM) | >229 µg/mL<sup>c</sup><br>(>10 µM) | >20 mg/Kg<sup>d</sup><br>(>400 µg/mL) |

<sup>a</sup>The Minimal Inhibitory Concentration 50% (MIC<sub>50</sub>) values measured after 10 h of histones treatment
are shown in µg/mL and their corresponding molarity value in parenthesis. Data is representative of
three independent experiments. <sup>b</sup>Lethal doses 50% (LD<sub>50</sub>) values in *G. mellonella* measured after 60 h
are indicated. <sup>c</sup>No bacterial growth inhibition was observed below the indicated concentration. <sup>d</sup>No
toxicity was found below 20 mg/Kg.

The biomass reduction of *P. aeruginosa* PAO1 planktonic cultures after 16 h of histone treatment at MIC_50_ (Table 1) was visualized and measured by Live and Dead staining (Fig. 1). A statistically significant decrease in bacteria cell number (58-70%) was shown in all samples treated with histone H1 subtypes compared to the control (Fig. 1A and B). Furthermore, some bacteria displayed red staining, indicating cell death (compromised membranes). Thus, we calculated the percentage of viable cells (green stained) present in each group (Fig. 1C) and found a statistically significant reduction in samples treated with H1.4 (31%), H1.0 (27%), and H1.2 (18%) compared to the control.

**Fig 1.**
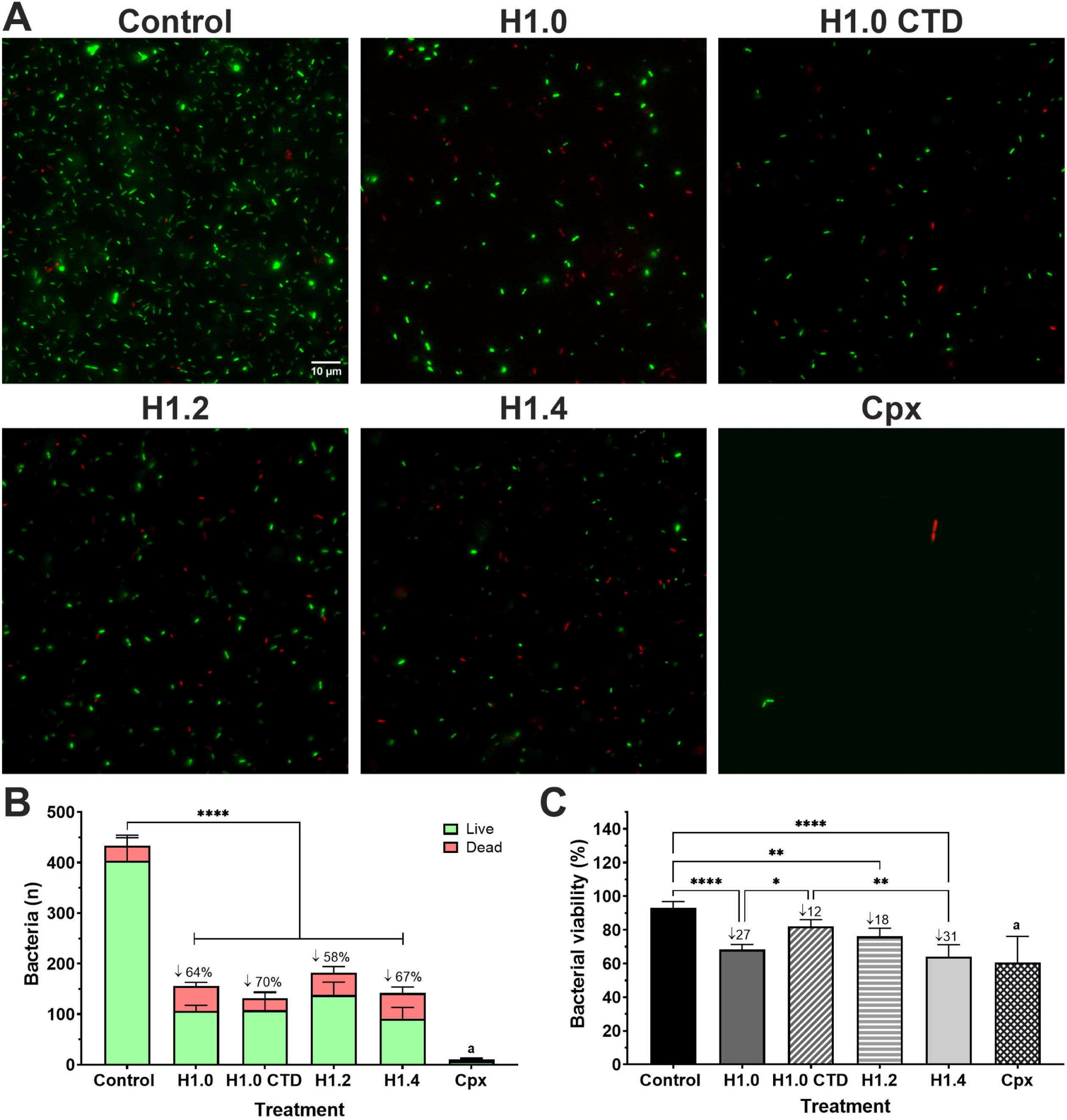
Bacterial viability assessment of *P. aeruginosa* PAO1 planktonic cultures after H1 histones treatment. (**A**) Live and dead staining using Syto 9 (green live cells) and Propidium iodide (red dead cells) dyes after 16 h of treatment. Histones were used at the MIC_50_ concentration of PAO1: H1.0 at 55 µg/mL, H1.0 C-ter 58 µg/mL, H1.2 at 111 µg/mL, and H1.4 at 46 µg/mL. Ciprofloxacin (Cpx) was used at 2 µg/mL. Fluorescence images were processed with Image J. The scale bar of 10 µm is consistent for all cases. (**B**) Quantification of live and dead bacteria by particle counting with Image J. Numbers after ↓ symbol indicate the percentage of reduction in bacteria total cell number compared to the untreated control. Error bars display mean and standard deviation from at least 4 replicates. Differences in the number of total cells among groups were analyzed by a one-way ANOVA analysis with Tukey’s multiple comparison test (∗∗∗∗, *p*-value <0.0001). Letter **a** represents a statistical difference of total cell number in the Cpx group compared to the control, H1.0 and H1.2 with a *p*-value <0.0001, and H1.0 CTD and H1.4 with a *p*-value <0.001. (**C**) Percentage of viability of bacterial cells ((Live bacteria/(Live bacteria + Dead bacteria))*100), where Live bacteria is the number of particles counted in the green channel (Syto 9 stained) and Dead bacteria the number of particles counted in the red channel (Propidium Iodide stained). Numbers after ↓ symbol indicate the percentage of reduction compared to the control. Error bars display mean and standard deviation from at least 4 replicates. Differences among groups were analyzed by a one-way ANOVA analysis with Tukey’s multiple comparison test (∗, *p*-value <0.05; ∗∗, *p*-value <0.01; ∗∗∗∗, *p*-value <0.0001). Letter **a** represents a statistical difference in the viability percentage of the Cpx group compared to the control with a *p*-value <0.0001, H1.0 CTD with a *p*-value <0.001, and H.12 with a *p*-value <0.05.

Consequently, we explored the potential use of histones H1.0 and H1.4 combined with antibiotics. We evaluated the synergy with Ciprofloxacin (Cpx) and Gentamicin (Gm) to inhibit the planktonic growth of PAO1. The results showed a synergistic effect only when histone H1.0 was combined with Cpx (Table 2), while the other combinations shown an additive effect.

**TABLE 2.** Evaluation of synergy between histones H1.0 and H1.4 with ciprofloxacin and gentamicin on *P. aeruginosa* PAO1 growth.

| Combination | FIC<sub>Histone</sub> | FIC<sub>Antibiotic</sub> | ΣFIC | Interaction |
| --- | --- | --- | --- | --- |
| H1.0 + Cpx | 0.25 | 0.25 | 0.5 | Synergy |
| H1.0 + Gm | 0.061 | 0.5 | 0.561 | Additivity |
| H1.4 + Cpx | 0.016 | 0.5 | 0.516 | Additivity |
| H1.4 + Gm | 0.012 | 0.5 | 0.512 | Additivity |

FIC: Fractional Inhibitory Concentration, Σ= Summatory, Cpx: Ciprofloxacin, Gm: Gentamicin.

### Histone and bacterial cells interaction

Aiming to elucidate the mechanism of action behind the antimicrobial activity of the H1 histones, we evaluated their direct interaction with *P. aeruginosa* PAO1 cells. Bacteria cultures were treated with histones for 30 min followed by a centrifugation step. Subsequently, we evaluated if histones were present in the supernatant or pellet fraction (Fig. 2). A precipitation control of histones without bacteria and a negative control of bacteria without histones were included. In Figure 2A, it is observed that irrespective of the histone, if bacteria were absent, western blot bands are observed both in pellet and supernatant. However, when mixed with bacteria, histones completely co-precipitate with them, suggesting a direct interaction of H1 histones with bacterial cells. Western blot bands were confirmed by immunoblot and post-transfer Coomassie staining (Fig. 2B), except in the case of H1.0 CTD due to unavailability of specific antibodies. Noteworthy, upon incubation with *P. aeruginosa* PAO1, H1 histones apparently suffered a partial cleavage, giving rise to smaller H1 forms present in the pellet.

**Fig 2.**
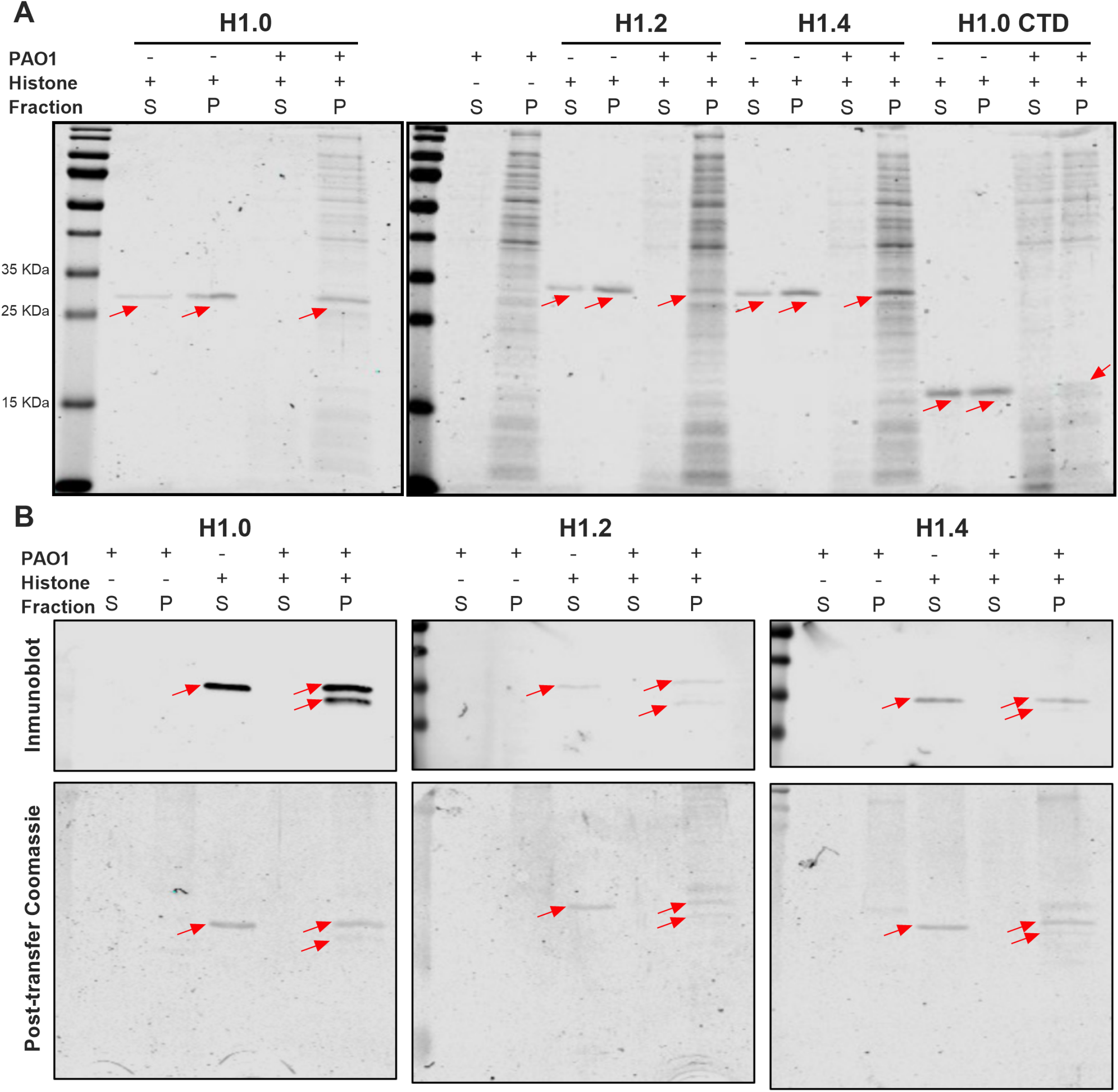
H1 histones precipitation assay after incubation of PAO1 planktonic cells. (**A**) Coomassie-stained SDS-PAGE gel of supernatant and pellet fraction from bacteria cultures, histones, and bacteria cultures treated with histones after centrifugation. Red arrows indicate the band corresponding to the respective histone subtype. (**B**) Inmunoblot and Post-transfer Coomassie staining of bands found in the supernatant and pellet fraction from bacteria cultures, histones, and bacteria cultures treated with histones after centrifugation. += Presence, - =Absence, S=Supernatant, P= Pellet.

Further, to assess if histone interactions caused any alteration to the bacterial cell membrane, cultures were treated with histones H1 at their *P. aeruginosa* PAO1 MIC_50_ concentration (Table 1) and stained with the specific dyes FM-464 and DAPI (Fig. 3). Some cells with membrane damage are observed in control samples after the centrifugation step (Fig 3. Control-zoomed images). However, very evident gaps in cell membranes were observed in all samples treated with histones (Fig. 3. Histones-zoomed images). Additionally, Histone H1.0 (Full length and CTD) and H1.4 caused non-homogenous staining among cells, with most showing a fainter membrane compared to the control sample. This effect is less markedly in samples treated with H1.2.

**Fig 3.**
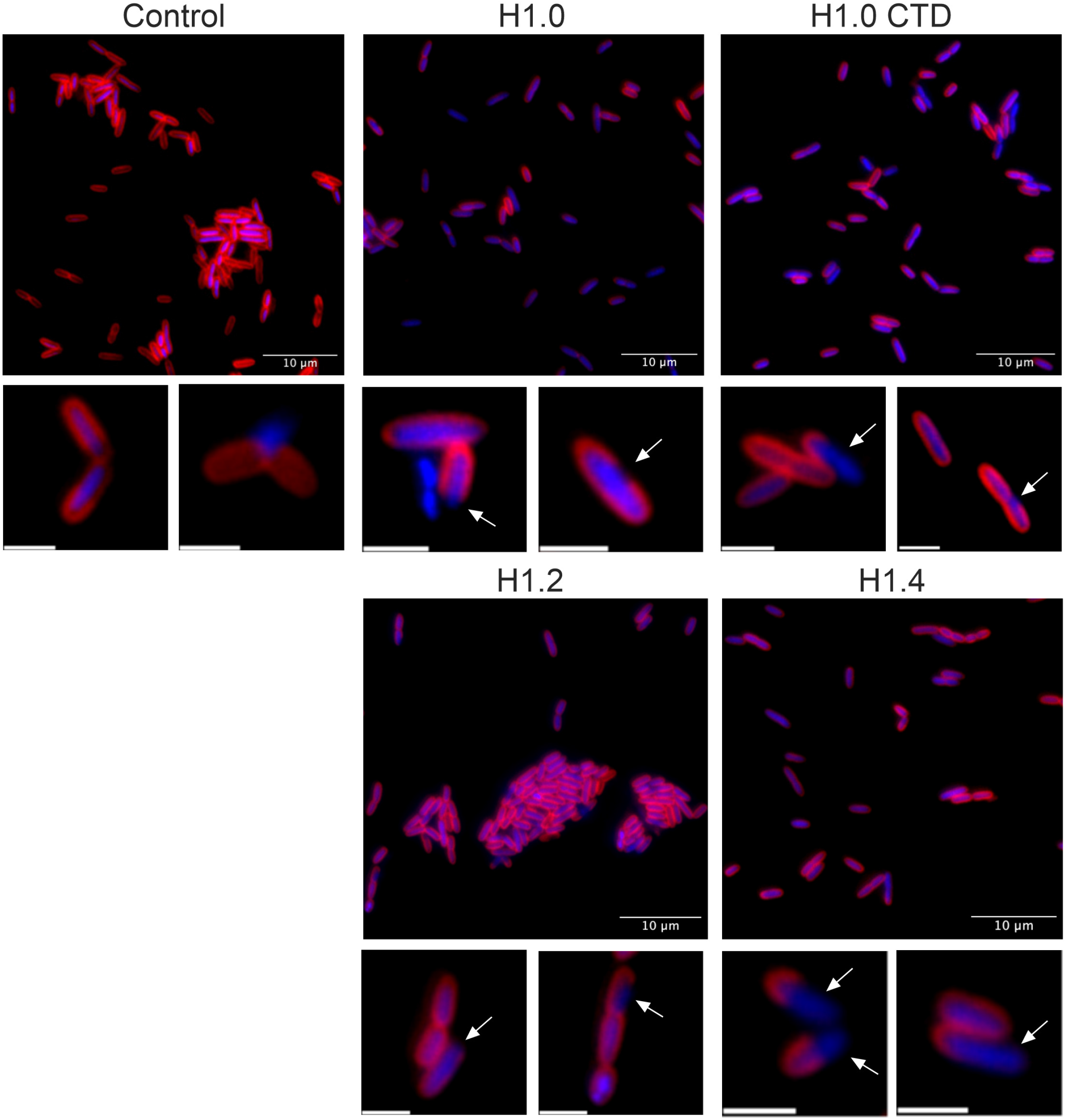
Assessment of membrane alterations of PAO1 cells after histones treatment. Bacteria were stained with FM-464 (red) and DAPI (blue) dyes to visualize the membrane and genetic material, respectively. Histones were used at the MIC_50_ of PAO1. Fluorescence images were processed with Image J software. Scale bars correspond to 10 µm in large images and 2 µm in zoomed images. White arrows point to gaps in cell membranes.

The differences observed in the alteration of bacterial membrane among histones subtypes could be explained by sequence and structural differences. Therefore, we analyzed several sequence-related properties of histone H1 subtypes, finding distinct features in the two subtypes with higher antimicrobial activity. Histone H1.0 has the highest positive charge density and the lowest hydropathicity, expressed as the GRAVY index, while H1.4 has the highest number of basic residues per protein molecule and highest net charge (Table 3).

**TABLE 3.** Protein characteristics of H1 subtypes.

| Subtype | Number of basic residues | Net charge | Positive charge density | GRAVY Index | Secondary structure prediction (%) |  |  |  |
| --- | --- | --- | --- | --- | --- | --- | --- | --- |
| | | | | | α-helix | Extended strand | Random coil | Ambiguous states |
| H1.0 | 62 | +53 | 0.32 | -1.07 | 35.57 | 4.12 | 50 | 10.31 |
| H1.2 | 62 | +55 | 0.29 | -0.68 | 32.39 | 2.35 | 55.87 | 9.39 |
| H1.4 | 66 | +59 | 0.30 | -0.78 | 38.81 | 2.28 | 46.12 | 12.79 |

Moreover, we studied the secondary structure of the recombinant proteins by bioinformatic analysis and experimental methods. Sequence predictions showed the highest values of α-helix in H1.4 followed by H1.0 and H1.2 (Table 3). The helical content and induction was studied by circular dichroism (Fig. S1). We calculated the ratio between the molar ellipticity in the minimum of the α-helix canonical spectrum at 222 nm, and the minimum of the entire spectrum in the random coil region, as an index of the proportion of α-helix present in each sample. The proteins were analyzed in aqueous solution, where the spectrum is clearly dominated by the random coil and in the presence of TFE, a known stabilizer of α-helix (19). We observed that, in contrast with the predictions, the molar ellipticity ratio was higher in H1.0 than in H1.4 and H1.2 in both conditions (Table 4).

**TABLE 4.** Circular dichroism of H1 subtypes.

| Solvent | Molar ellipticity ratio<sup>a</sup> |  |  |
| --- | --- | --- | --- |
|  | H1.0 | H1.2 | H1.4 |
| PBS | 0.55 | 0.43 | 0.42 |
| TFE 20% | 0.63 | 0.56 | 0.57 |

<sup>a</sup>Molar ellipticity ratio was calculated as the value at 222 nm referred to the minimum of the spectrum. TFE= Trifluoroethanol.

To evaluate the existence of a correlation in the variation of the parameters of positive charge, hydropathicity and, α-helix content and the differences observed in the activity of histone H1 subtypes we performed simple linear regressions. As a result, any correlation with the antimicrobial activity of histones was found (reduction of planktonic cells after treatment). However, there is a positive correlation between the proportion of α-helix content induced in TFE and the antibiofilm activity of the histone H1 subtype against static biofilms of PAO1 (r^2^=1, *p*-value=0.0043). Unfortunately, the low number of evaluated proteins made not possible the performance of a multiple linear regression that evaluates simultaneously the effect of the parameters.

### Histone H1 antibiofilm efficacy

Recognizing the significance of biofilm related infections, we investigated the efficacy of histones against *P. aeruginosa* PAO1 biofilms. Initially, we assessed the reduction in biofilm biomass after treatment with H1 histones using a static biofilm model (96-well plate) and the crystal violet biomass staining. As shown in Figure 4A, all histone-treated samples exhibited some degree of biomass reduction, with more pronounced effects observed when using H1.0 (28%, statistically significant), followed by H1.4 (22%) and H1.2 (20%).

**Fig 4.**
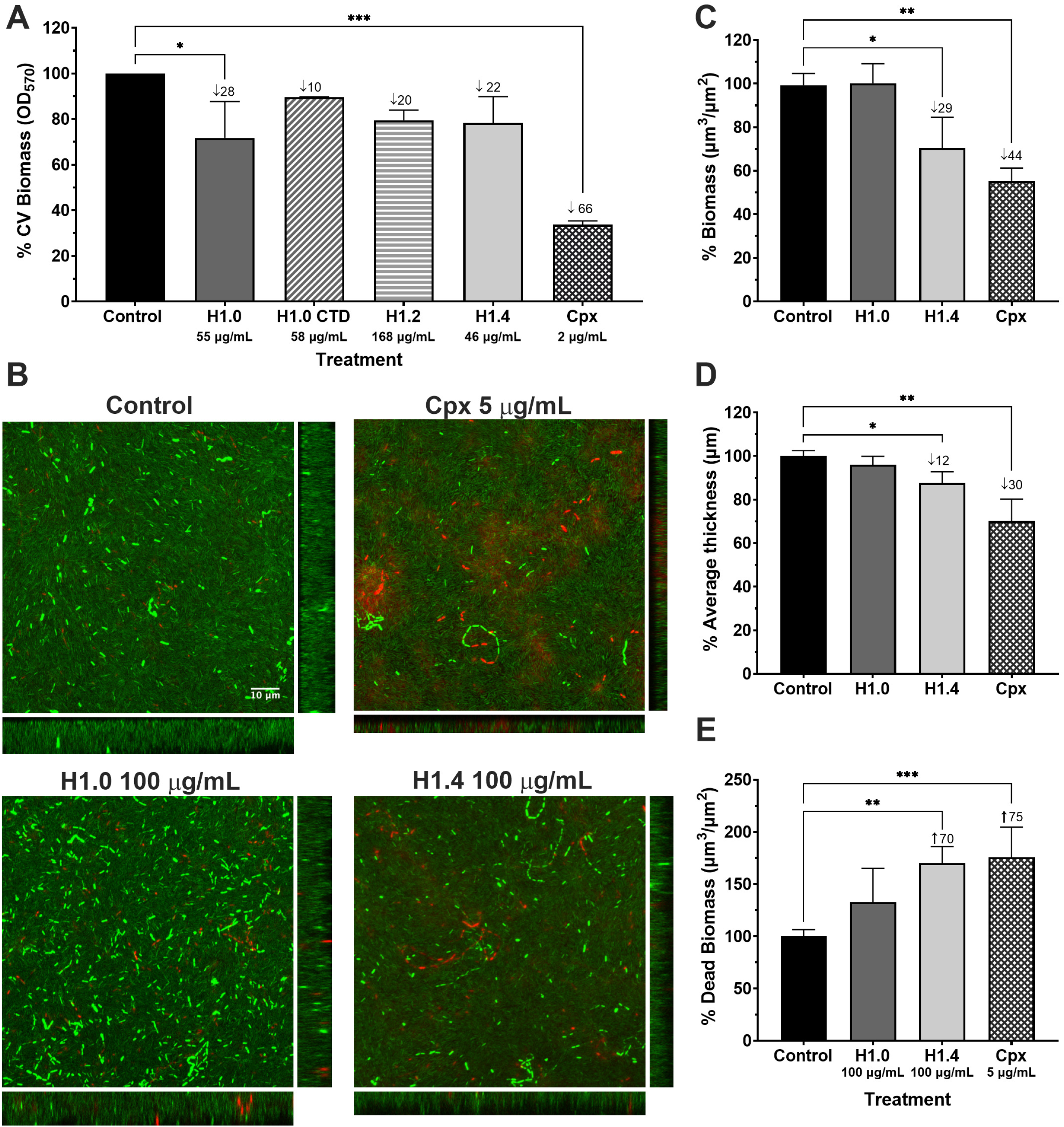
Antibiofilm activity of histones against PAO1 biofilms. (**A**) Biomass reduction of static biofilms after histone treatment. Error bars display mean and standard deviation of at least three replicates. Data differences with respect to the control group were analyzed by a one-way ANOVA analysis with Dunnett’s multiple comparison test (∗, *p*-value <0.05; ∗∗∗, *p*-value <0.001. (**B**) Confocal microscopy visualization of continuous flow biofilms after treatment with H1.0 and H1.4. Biofilms were stained with Syto 9 and Propidium Iodide to dye live cells in green and dead cells in red. Z-stacks from confocal images with their corresponding orthogonal views are displayed. The scale bar of 10 μm is consistent across all images. Confocal images analysis of the (**C**) Percentage reduction in biomass, (**D**) Percentage reduction in Average thickness percentage, and (**E**) Percentage increase in dead Biomass after treatment of flow biofilms with H1.0 and H1.4. Numbers after the ↓ symbol indicate the percentage of reduction compared to the control, while numbers after the ↑ symbol indicate the percentage of increase compared to the control. Error bars display mean and standard deviation of at least three replicates. Data differences with respect to the control group were analyzed by a one-way ANOVA analysis with Šidák’s multiple comparison test (∗*p*-value <0.05; ∗∗*p*-value <0.01; ∗∗∗*p*-value <0.001). Cpx= Ciprofloxacin.

We selected H1.0 and H1.4 subtypes to test their antibiofilm *in vitro* efficacy in a continuous flow chamber biofilm assay, a model that better resembles *in vivo* biofilm infections. PAO1 biofilms treated with 100 μg/mL of each H1 subtype were died with the Live and Dead staining and imaged by confocal microscopy (Fig. 4B). As seen in Figure 4B, the orthogonal views show a significant decrease in bacterial biomass in the biofilm treated with histone H1.4 compared to the untreated control. Quantification confirmed a significant reduction in PAO1 biofilm biomass (29%) and average thickness (12%) after H1.4 treatment (Fig. 4C and D). Moreover, the dead biomass (red staining) significantly increased in the biofilm treated with H1.4 (Fig. 4E). No significant effect in the evaluated parameters was observed after H1.0 treatment. The findings obtained under this grown condition are particularly noteworthy because the resulting biofilms are highly resistant to antibiotics as seen with the Cpx treatment as positive control.

### *In vivo* H1.0 and H1.4 toxicity assessment and antimicrobial efficacy

Next, we used the *G. mellonella* larvae infection model to evaluate the antimicrobial potential of histone H1. No mortality or morbidity parameters (myelinization, activity reduction and absence of cocoon) (20) associated with toxic effects were observed in *G. mellonella* larvae for up to 60 h post-injection with H1.0 and H1.4 histones in a range of 1.25-20 mg/Kg (25 µg/mL-400 µg/mL), indicating low toxicity.

Once histones toxicity was discarded, the use of histones H1.0 and H1.4 as treatments for recovering infected larvae with PAO1 was evaluated. 1 h post-infection, larvae were treated with histones (100 μg/mL) alone and in combination with Cpx (Fig. 5). Larvae survival was increased and extended when infected *G. mellonella* were treated with histones compared to those treated with PBS (negative control). Specifically, the median survival was significantly extended from 16 h in PBS treated group to 18 h and 20 h in the histone H1.0 and H1.4 treated group, respectively. However, when comparing larvae treated with Cpx 1.5 mg/Kg versus those treated with the combined therapy, no significant improvement was observed.

**Fig 5.**
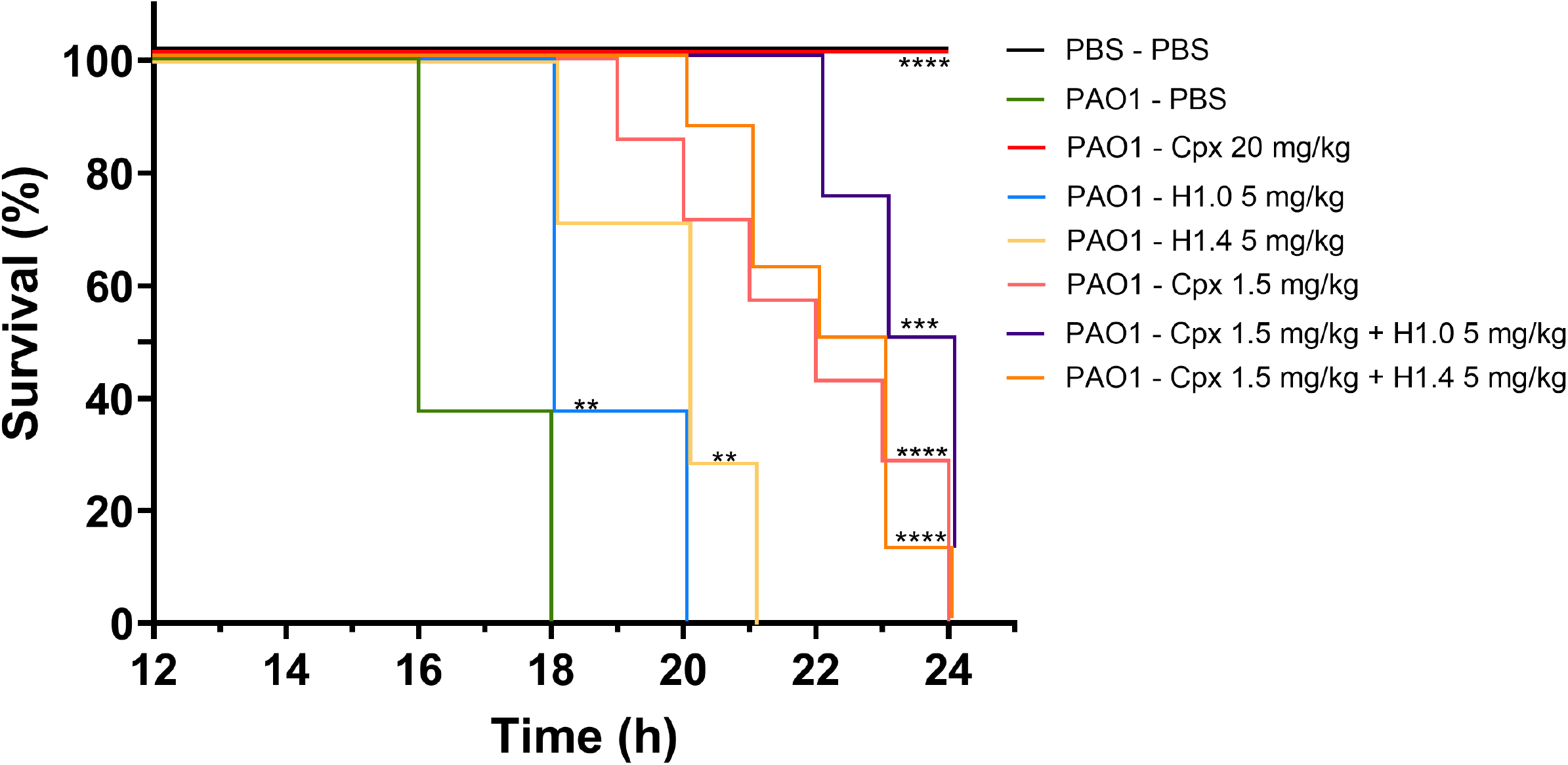
*In vivo* evaluation of H1 histones activity against *G. mellonella* infected with PAO1. Kaplan-Meier survival curves of *G. mellonella* larvae infected with PAO1 and treated with histones alone or in combination with Ciprofloxacin (Cpx). Control larvae were injected with PBS. Larval mortality was monitored for 16-24 h post-injection with observations conducted hourly. Significant differences in treatment efficacy compared to the PAO1-PBS treated group were assessed using a log-rank test (∗, *p*-value <0.05; ∗∗, *p*-value <0.01; ∗∗∗, *p*-value <0.001; ∗∗∗∗, *p*-value). The results depicted in this figure represent an experiment replicated three times with consistent outcomes, each condition involving 8 larvae.

## Discussion

Histones are ubiquitous across different tissues and their antimicrobial activity had been reported in multiple species (10). Moreover, some peptide fragments derived from histones cleavage are considered AMPs (21). For instance, H1 histones secreted by human colonic epithelial cells possess antimicrobial activity (15, 22), as well as the peptides derived from the NTD of H1 histone from Atlantic salmon and the CTD of H1 histone from rainbow trout (known as Oncorhyncin II) (23, 24).

In this work, we reported the ability of three human H1 subtypes (H1.0, H1.2, and H1.4) to inhibit bacterial proliferation and induce cell death in three well-known opportunistic pathogens: *P. aeruginosa*, *S. aureus* and *E. coli* (Table 1 and Fig. 1). Our findings support the study by Jacobsen *et al*. (2005), where the efficacy of recombinant human histone H1.2 was demonstrated against bacteria found in human wound infections (25). Similarly, Jodoin & Hincke (2018) reported that histone H1 from chicken (named H5 in this animal) has antimicrobial activity against Gram-negative and Gram-positive bacteria (17).

Overall, our results (Table 1) demonstrated that regardless of the H1 histone subtype used, a lower concentration of the protein is required to inhibit Gram-negative bacteria (*P. aeruginosa* PAO1 and *E. coli* CFT073). This observation is consistent with the affinity of H1 for bacterial lipopolysaccharide (LPS), the major component of the outer membrane of Gram-negative bacteria (26). Our study confirms an interaction between H1 histones and *P. aeruginosa* PAO1 cells upon mixed incubation (Fig. 2). Moreover, the fragmentation of all three histone subtypes in the precipitate fraction, alongside bacteria, suggests that bacterial proteases may degrade the histones, and their antimicrobial properties could be attributed to AMPs characteristics.

The mechanism underlying histone-mediated bacteria killing remains unclear, but two primary options have been proposed: membrane disruption or cytoplasmatic targeting. H1 from the liver of Atlantic salmon causes morphological alterations that directly damage the cell surface (27). Likewise, H1 from chickens caused pore formation and content liking in *P. aeruginosa* cells (17). Conversely, Onconrhyncin II is unable to form stable channels in the membrane, suggesting an antimicrobial activity resultant from intracellular processes (24). In this study, bacterial cells treated with histone H1 subtypes displayed clear membrane gaps, indicating bacterial killing by alteration of membrane integrity (Fig. 3). Furthermore, we hypothesized that their presence in the bacterial membrane hindered FM-464 dye incorporation, resulting in less intense staining in histone treated samples compared to the untreated control. This effect was more pronounced in the case of H1.0 and H1.4, suggesting a higher affinity of these histones with *P. aeruginosa* PAO1 membrane.

Several factors can influence how an AMP causes membrane alterations, including propensity for peptide self-assembly, net charge, and amphipathicity (21, 28). These characteristics can be found in peptides with an amphipathic α-helix structure (30, 31). We compared the positive charges, hydropathicity and α-helix content of H1.0, H1.2, and H1.4 proteins (Table 3 and 4). Overall, H1.0 and H1.4, the subtypes with higher antimicrobial and antibiofilm activity, exhibited higher net charge, lower gravy index (more hydrophilic) and higher α-helix content. Simple linear regression models showed a correlation between α-helix content induced in TFE and the antibiofilm activity of H1 subtypes against *P. aeruginosa* PAO1 static biofilms. However, the structural properties of the three subtypes are very similar and more robust statistical analysis need to be conducted.

As not all the antimicrobial molecules had antibiofilm activity, we decided to test H1 histones’ efficacy against *P. aeruginosa* PAO1 biofilms (Fig. 4). The urgent need for alternative molecules that can eradicate biofilms, specially from multi-drug resistant bacteria, has led to the identification of several AMPs with antibiofilm properties (see the Biofilm-active AMPs database at http://www.baamps.it/) (32). However, as far as we are aware, the ability of histones for biofilm disruption has been understudied. Jacobsen *et al.* showed a reduction of ∼60% of bacterial burden in an *in vivo* rat burn model infected with *P. aeruginosa* after treatment with H1.2 protein (25). Likewise, the work from Jodoin & Hincke (2018) reported that the Mimimum Biofilm Erradication Concentration (MBEC) value of H1 from chicken against *P. aeruginosa* biofilms formed under static conditions is >128 µg/mL (17). Although, the methodology used in our study is diferent, our data pointed to a similar MBEC value.

We used two different *in vitro* models widely employed for biofilm studies. The microtiter plate, a static model used, easy to use and high throughput; and the continuous flow chamber assay, a dynamic model in which the resulting biofilms are structurally similar to those present in natural infections (33). Flow conditions stimulate the production of robust ECM, which, in the case of *P. aeruginosa* PAO1 biofilms, contain alginate and extracellular DNA, two highly negatively charged molecules that can trap positively charged agents (34). Thus, we expected this scenario to be more challenging for histone treatment efficacy. We found higher antibiofilm activity in H1.0 in the static model compared to the other H1 subtypes (Fig. 4A). However, in the dynamic biofilm model, H1.4 has better results than H1.0 (Fig. 4B and D). H1.0 has a higher density of positive charges, lower gravy index (more hydrophilic) and, higher formation of α-helix in TFE compared to H1.4 (Table 3). Similarly, H1.0 CTD, the protein region with more positive charges and less hydrophobicity (12, 35) showed reduced activity against *P. aeruginosa* PAO1 biofilms (Fig. 4A) when compared to H1.0 (full length). This could indicate that these parameters are influencing the antibiofilm activity of histones, being their effect more markedly when biofilms are formed in dynamic conditions.

It is important to consider that the biofilm reduction assessed by the static crystal violet assay quantifies both cell and ECM biomass, while Live and Dead staining only assess differences in cell biomass. Nevertheless, we found the almost 30% reduction in cell biomass obtained after H1.4 treatment in the flow biofilm assay to be highly significant (Fig. 4B and C). Considering the reduction observed in the average biofilm thickness (Fig. 4D) and the increase in the biomass stained with propidium iodide (Fig. 4E), we proposed that the antibiofilm activity of the histone H1.4 is associated with its capacity to induce bacterial cell death.

Finally, the antimicrobial activity of H1 subtypes against *P. aeruginosa* PAO1 obtained in this study was validated in an *in vitro* animal model. It is known that the use of histones as therapeutic agents could lead to adverse effects. For instance, extracellular histones are damage-associated molecular patterns, able to induce chemokine release and cytotoxicity (10). They can induce thrombin generation and platelet aggregation, promoting thrombosis and thrombocytopenia (21). Moreover, histone interaction with cell membranes can activate adverse inflammatory responses that worsen several diseases (21)

Regarding histone H1, Onconrhyncin II and H1 from chicken do not have significant hemolysis (17, 24). Similarly, human recombinant H1.2 showed very low hemolytic activity (2%) at 250 µg/mL, but cytotoxicity against human keratinocytes at low concentrations was observed (25). In this study, we did not observe any sign of toxicity in *G. mellonella* larvae injected with histone H1.0 and H1.4 up to 20 mg/Kg (400 µg/mL). Furthermore, we found an increase of the median survival rate after treatment of *G. mellonella* infected with *P. aeruginosa* PAO1 (Fig. 5). We are aware that further toxicity and efficacy studies involving human cells and mammals must be conducted, but the positive results obtained against this aggressive bacterium are promising. In addition, the discovery that polysialic acid can reduce the H1-mediated cytotoxicity in eukaryotic cells without alter its antimicrobial activity creates a new alternative in biomedical applications (9).

There is still a long road and several limitations to overcome before the implementation of histone-derived AMPs, and AMPs in general, be consolidated in clinical context (36, 37). However, day-to-day new strategies are proposed (38–40). For instance, the concomitant use of histones with AMPs with different action modes has been reported efficient (41). Similarly, AMPs combined with antibiotics had proven enhanced antibacterial activity (8). In this work, we found synergy between Cpx and H1.0 *in vitro* (Table 2), but any significant difference in median survival rate of *G. mellonella* larvae was observed with the combined treatment compared with the antibiotic alone (Fig. 5). More studies regarding the action mode of H1 histones could help to elucidate these findings.

In conclusion, we have demonstrated that H1.0, H1.2 and H1.4 have antimicrobial activity against *P. aeruginosa* PAO1, likely attributable to membrane disruption. H1.0 and H1.4 were the subtypes that exhibit higher antimicrobial activity with lesser concentration required. H1.4 had higher antibiofilm potential and better survival outcomes *in vivo* in an acute infection model of *P. aeruginosa* PAO1. Although we did not validate a synergistic effect of H1.0 and H1.4 with the antibiotics Ciprofloxacin and Gentamicin, none of the histone subtypes caused toxicity to *G. mellonella* larvae.

## Methods

### Histones expression and purification

Histones H1.2 was cloned into pET21a and expressed in *Escherichia coli* BL21(DE3) (Merk Millipore) while H1.0 and its C-terminal domain (CTD) were cloned into pQE60 and expressed in *E. coli* M15 (42). H1.4 was cloned in pET21a using the forward primer (*Nde*I) 5’-TAATATCATATGTCCGAGACTGCGCCTGCC-3’ and the reverse primer (*Xho*I) 5’-ATATATCTCGAGCTTTTTCTTGGCTGCCGCCT-3. Expression was carried out in *E. coli* BL21 (DE3).

All proteins contained a His-tag at the C-terminal protein end. Recombinant proteins were expressed and purified by metal-affinity chromatography as previously described (42). Before their use, histones were thawed, vortexed, and sonicated (3 cycles of 20 s) using an ultrasonic processor UP50H (Hielscher ultrasonics).

### Sequence analysis and predictions

The sequences of H1.0, H1.2, and H1.4 were used to compare the positive charge density and the predicted secondary structure of the analyzed subtypes. Positive charge density of H1.0 (P07305), H1.2 (P16403), and H1.4 (P10412) was calculated as the quotient between the number of basic residues and the protein length. The GRAVY index was calculated by the sum of hydropathy values of all amino acids divided by the protein length. Consensus secondary structure predictions were obtained at the Network Protein Sequence @nalysis (NPS@) server (https://prabi.ibcp.fr/htm/site/web/app.php/home).

### Circular dichroism

Secondary structure of H1.0, H1.2, and H1.4 was analyzed in aqueous solution (PBS) and in the presence of 20% 2,2,2 trifluoroethanol (TFE). Proteins were resuspended in the desired solvent at a final concentration of 230 ng/µL. Measurements were recorded in a J-815 spectropolarimeter (JASCO) in a continuous scanning mode, in the 190-250 nm range at a scanning speed of 200 nm/min with a bandwidth of 1 nm. Five accumulations were taken of each spectrum. Molar ellipticity was obtained by normalizing CD units by the protein concentration and the number of residues.

### Bacterial strains and growing conditions

Three different bacterial strains were used in this study: *Pseudomonas aeruginosa* PAO1 strain (CECT 4122 – ATCC 15692), referred to as PAO1, *Staphylococcus aureus* (CECT 86 – ATCC 12600), and an uropathogenic *E. coli* CFT073 strain (ATCC 700928). Bacteria were retrieved from a −80°C stock and cultured on Luria-Bertani media (Scharlau) for PAO1 and *E. coli* CFT073, and Tryptic Soy agar/broth (TSA/TSB) (Scharlau) for *S. aureus*. Overnight cultures were made by incubation at 37°C and 200 rpm. Unless specified, bacteria inoculum for each experiment were prepared by incubating 100 μL of overnight cultures in fresh TSB media until reaching the initial exponential log phase (OD_550 nm_ ≈ 0.2-0.3).

### Antibacterial activity against planktonic bacteria

100 μL of inoculum from all three bacterial strains were plated in a 96-well microtiter plate (Corning) filled with different histone protein concentrations. TSB media was used as a negative control. Following a previously described method (18), the microtiter plate was incubated at 37 °C in a SPARK Multimode microplate reader (Tecan) with 150 rpm shaking. Bacterial growth was monitored for 10 h with readings taken every 15 min.

The minimal inhibitory concentration 50% (MIC_50_) was defined as the compound concentration that reduces bacterial growth, determined as the OD_550_, by 50%.

### Fluorescent microscopy viability test analysis

Aliquots of 500 μL from a *P. aeruginosa* PAO1 inoculum were treated with histones at a concentration equal to their MIC_50_ value of each subtype (See Table 1). Samples treated with Ciprofloxacin (Cpx) at 2 µg/mL and media were included as controls. After 16 h of incubation at 37°C, 100 μL of the samples were centrifuged at 6000 rpm for 5 min. Pellets were resuspended in 25 µL of PBS (Fisher Scientific S.L.) and stained with Syto 9 and Propidium iodide (Live/Dead BacLight Bacterial Viability Kit, Thermo Fisher Scientific) following the manufacturer’s instructions. Samples were visualized using a Nikon inverted fluorescent microscope ECLIPSE Ti–S/L100 (Nikon, Japan) coupled with a DS-Qi2 Nikon camera (Nikon, Japan). Images were processed and quantified by analysis of particles with Image J software. The percentage of viability was calculated according to the following formula:

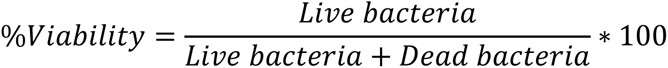

where Live bacteria is the number of particles counted in the green channel (Syto 9 stained) and Dead bacteria the number of particles counted in the red channel (Propidium Iodide stained).

### Antibiotic-histones synergy testing by checkerboard assay

The synergy of histones H1.0 and H1.4 in combination with ciprofloxacin (Cpx) and gentamicin (Gm) was determined by the standard broth microdilution assay as described previously (43). Briefly, *P. aeruginosa* PAO1 inoculum was added in a 96-well microtiter plate with increasing concentrations of histone on one axis and increasing concentrations of antibiotic on the other. Plates were incubated at 37 °C in a SPARK Multimode microplate reader with 150 rpm shaking for 16 h. The effect of the antimicrobial combination was defined using the lowest Fractional inhibitory concentration (FIC) index (44). The formula employed for FIC calculation (43) and its interpretation (45) were as previously described:

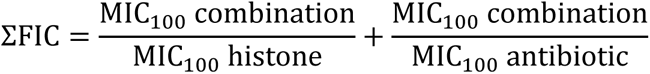

Where ΣFIC ≤0.5 is defined as synergy, ΣFIC >0.5 and <4 is additivity/indifference and, ΣFIC >4 is antagonism.

### Histones and bacterial cells interaction by precipitation experiments

Aliquots of 200 μL of a *P. aeruginosa* PAO1 inoculum were treated with a concentration of 10 μg/mL of each histone subtype. After 30 min of incubation at 37°C and 300 rpm in a thermomixer (Eppendorf), samples were centrifuged at 6000 rpm for 5 min, and supernatant and pellet were collected. Histones added to TSB were used as precipitation control.

Subsequently, supernatant and pellet samples were boiled (5 min at 95°C) and exposed to SDS-PAGE (14%), followed by Coomassie staining or immunoblotting with H1 variant-specific antibodies. For immunoblotting, protein extracts were transferred to PVDF membrane, blocked with 5% non-fat milk for 1 h, and incubated overnight at 4°C with the primary antibodies anti-H1.0 (Millipore 05-629l), anti-H1.2 (Abcam ab4086), and anti-H1.4 (Invitrogen 702876). No specific antibodies against the C-terminal region of H1.0 were available. Next, samples were incubated with secondary antibodies conjugated to fluorescence (IRDye 680 goat anti-rabbit IgG or IRDye 800 goat anti-mouse IgG, Li-Cor) for 1 h at room temperature. Bands were visualized using an Odyssey Infrared Imaging System (Li-Cor). Coomassie staining was used as a loading control.

### Bacterial membrane integrity after H1 incubation

Aliquots of 200 μL of a *P. aeruginosa* PAO1 inoculum were treated with histones H1.0, H1.0 CTD, H1.2 and H1.4 at a concentration of 55 µg/mL, 58 µg/mL, 111 µg/mL, and 46 µg/mL, respectively, which correspond to their MIC_50_ value (Table 1). After 30 min of incubation at 37°C and 300 rpm in a thermomixer (Eppendorf), samples were centrifuged at 6000 rpm for 5 min. Pellets were resuspended in 50 µL of PBS with DAPI (Invitrogen) at 100 μg/mL and FM® 4-64 (Invitrogen) at 40 μg/mL for 15 min. Samples were dropped on a slide pre-covered with agarose 1% (w/v) as previously described (46) and visualized with a Nikon inverted fluorescent microscope ECLIPSE Ti–S/L100 (Nikon, Japan) coupled with a DS-Qi2 Nikon camera (Nikon, Japan). Images were processed with Image J software.

### Antibacterial efficacy against biofilms

For static biofilm analysis, an overnight culture from *P. aeruginosa* PAO1 was diluted to OD_550 nm_ = 0.1 in TSB supplemented with 0.2% glucose (Fisher Scientific S.L.), added in a 96-well microtiter plate, and incubated at 37 °C. After 72 h, wells were washed three times with PBS and treated with the different H1 histones at PAO1 MIC_50_ (Table 1). TBS and Cpx at 2 µg/mL-treated wells were used as controls. 6 h after treatment, three PBS washes were carried out, and wells were fixed with methanol (Fisher Scientific) for 15 min. Then 1% (w/v) Crystal violet (Merck Life Science) was added for 5 min followed by a distaining step with 30% acetic acid (Scharlau). Biofilm biomass was determined by measuring the absorbance (OD_570_) using a Microplate spectrophotometer Benchmark Plus (Bio-Rad, USA).

The antibiofilm efficacy of histones H1.0 and H1.4 was also tested in a continuous PAO1 flow biofilm assay as previously described (18). Briefly, an overnight culture diluted at OD_550 nm_ =1 was inoculated into a flow chamber and allowed to attach for 2 h. Then, fresh TSB + 0.2% glucose was continuously pumped (42 µL/min) through the flow chamber for 72 h until a mature biofilm was obtained. The mature biofilms were treated with histones at 100 µg/mL and left to act for 6 h. Treatment with TBS and Cpx at 5 µg/mL were used as controls. Finally, biofilms were stained with Syto 9 and Propidium iodide according to the manufacturer’s instructions and visualized with a Zeiss LSM 800 confocal laser scanning microscope (CLSM). Image processing was performed with Image J software, and measurements of biofilm biomass (green channel), average thickness (green channel), and dead biomass (red channel) were obtained using FIJI and COMSTAT2 plugins (47).

### Antibacterial efficacy and toxicology in a *Galleria mellonella* infection model

Maintenance and injection of *G. mellonella* larvae were carried out as previously reported by our laboratory (48). Larvae selected in all experiments had a weight range of 175-250 mg.

First, the toxicity of H1.0 and H1.4 histones was evaluated by injecting 5 different concentrations into the larvae: 1.25 mg/kg, 2.5 mg/kg, 5 mg/kg, 10 mg/kg, and 20 mg/kg. Considering that 2.5 mg/kg is equivalent to 50 µg/mL, the evaluated concentration range comprises ∼0.5 x MIC_50_ to ∼8 x MIC_50_ of *P. aeruginosa* PAO1 to H1 histones subtype (Table 1). PBS treated larvae were included as control. Additionally, larvae injected with PBS were treated after 1 h with histones at 20 mg/kg to eliminate mortality due to consecutive injuries from the injection process. The experiment was conducted twice and included a total of 5 larvae per group each time. Larvae were monitored for 60 h.

Then, the antimicrobial efficacy of histones against *P. aeruginosa* PAO1 infection in *G. mellonella* was tested. Larvae were infected with *P. aeruginosa* PAO1 (5-50 CFUs/larvae) and 1 h after, 5 mg/kg of histone H1.0 and H1.4 were injected alone and in combination with 1.5 mg/kg of Cpx. Infected larvae were also treated with PBS as a negative treatment control, Cpx 20 mg/kg as positive control of treatment and, Cpx 1.5 mg/kg as a control of synergy effect between antibiotic and histones. Moreover, a group of larvae injected with PBS at the infection and treatment step were included as control of double injection. The experiment was conducted three times with 8 larvae per group. Larvae were monitored from 16 h-24 h post-infection, with observations every hour.

### Statistical analysis

All data were statistically analyzed using GraphPad Prism version 10.00 (GraphPad Software, USA). Comparison of means among groups were performed by One-way ANOVA tests. Correlations among structural parameters and the antimicrobial and antibiofilm effect of histone H1 subtypes were evaluated by simple linear regressions. Comparison of Kaplan-Meier survival curves were made by Log-rank tests. A *p*-value <0.05 was considered statistically significant.

## Acknowledgements

This study was partially supported by grants PID2021-125801OB-100, PLEC2022-009356 and PDC2022-133577-I00 to ET, PID2020-112783GB-C21 to A.J and PID2020-112783GB-C22 to A.R., funded by Spanish Ministry of Science and Innovation MCIN/AEI/10.13039/501100011033 and “ERDF A way of making Europe”, the CERCA programme and *AGAUR-Generalitat de Catalunya* (2021SGR01545), the European Regional Development Fund (FEDER) and Catalan Cystic Fibrosis association. The project that gave rise to these results received the support of a fellowship from “la Caixa” Foundation (ID 100010434). The fellowship code is “LCF/BQ/DI20/11780040”.

The funders had no role in study design, data collection and interpretation.

